# High-performance proteomics at any chromatographic flow rate

**DOI:** 10.1101/2025.04.10.648108

**Authors:** Giorgi Tsiklauri, Runsheng Zheng, Nicole Kabella, Polina Prokofeva, Christopher Pynn, Bernhard Kuster

## Abstract

Current applications of mass spectrometry-based proteomics range from single cell to body fluid analysis that come with very different demands regarding sensitivity or sample throughput. Additionally, the vast molecular complexity of proteomes and the massive dynamic range of protein concentrations in these biological systems require very high-performance chromatographic separations in tandem with the high speed and sensitivity afforded by mass spectrometer. In this study, we focussed on the chromatographic angle and, more specifically, systematically evaluated proteome analysis performance across a wide range of chromatographic flow rates (0.3 – 50 μL/min) and associated column diameters using a Vanquish Neo UHPLC coupled online to a Q Exactive HF-X mass spectrometer. Serial dilutions of HeLa cell line digests were used for benchmarking and total analysis time from injection-to-injection was intentionally fixed at 60 minutes (24 samples per day). The three key messages of the study are that i) all chromatographic flow rates are suitable for high-quality proteome analysis, ii) capLC (1.5 μL/min) is a very robust, sensitive and quantitative alternative to nanoLC for many applications and iii) showcased proteome, phosphoproteome and drug proteome data provide sound empirical guidance for laboratories in selecting appropriate chromatographic flow rates and column diameters for their specific application.

## INTRODUCTION

Nano-flow (<1 µL/min) LC-MS/MS has been the cornerstone of proteome analysis technology for nearly 30 years, due to its exceptional sensitivity when coupled to electrospray ionization mass spectrometry, a crucial requirement in the early days of proteomics. Over the past decade, significant advancements in mass spectrometric instrumentation have massively enhanced both sensitivity and speed of proteome analysis. Progress in the former has led to impressive results when studying biological systems from which only very small quantities of protein are available. The latter has increased the opportunity for applying proteome analysis in a screening mode exemplified, for instance, by the analysis of clinical patient cohorts, protein-protein or protein-drug interactions. In many of these scenarios, sample throughput, robustness, and reproducibility of LC-MS/MS systems are just as, or even more important than sensitivity. These requirements have sparked renewed interest in liquid chromatography separation at higher flow rates [1–3] because all parameters except sensitivity often see marked improvements when higher flow rate systems are employed. These systems are often more suitable for high-throughput studies where exceptional sensitivity may not be of paramount importance. Higher flow alternatives to nano-flow separations have been discussed in detail in a recent review [4]. Briefly, an elegant study published in 2018 described a μLC-MS/MS setup (1 mm internal diameter (i.d.) × 25 cm column, 68 µL/min) identifying approximately 2,800 proteins in 1 hour from 2 µg of HeLa tryptic peptides using a Q-Exactive Orbitrap mass spectrometer, demonstrating the feasibility of μLC–MS/MS with reasonable sample requirements [5]. Bian et al. slightly modified the approach and showed that the proteomic depth of nLC-MS/MS could be matched by microflow LC-MS/MS (μLC-MS/MS) when using 5-10 times more peptide amounts [6]. The same authors later demonstrated that such a system is extremely robust (>38,000 samples in two years, >14,000 proteomic analyses on a single column) [7]. Consequently, μLC-MS/MS has been used in a diverse range of proteomic studies, highlighting its versatility and reliability [8, 9].

Capillary flow LC-MS/MS (capLC-MS/MS) has also emerged as a promising alternative to nLC-MS/MS but its utility has not been as well explored. Early work by Tao et al. demonstrated its potential by identifying 1,692 proteins from rat brain tissue using a 300 µm i.d. column operated at 5 µL/min [10]. In 2018 the Ralser laboratory optimized a 300 µm i.d. column running at 3-10 µL/min and using SWATH-MS to quantify 4,000 human and 1,750 yeast proteins in under one hour [11]. Similarly, Bruderer et al. explored a 0.3 mm i.d. column at 5 µL/min with DIA and managed to analyze 31 human plasma proteomes in 24 hours and analyzed 1,508 human plasma samples from a nutritional intervention study cohort [12]. Very recently, the ProCan team in Australia showcased a 300 μm i.d. column setup running at 5 μL/min for high-throughput proteomics and phosphoproteomics of rat tissues using a ZenoTOF instrument reaching 2,600 protein identifications in 30 min gradient time from 400ng rat brain peptides [13]. Also recently, Sui et al. reported results from comparing μLC-MS/MS (1 mm i.d. 50 µL/min) to capLC-MS/MS (150 µm i.d. 1 µL/min) and concluded that it was suitable for high-throughput analysis of clinical samples with limited available material [14].

While there are several comparative studies showing that reducing flow rate enhances sensitivity and higher-flow rate offers higher throughput and robustness, no single study has compared the performance of nLC-, capLC- and μLC-MS/MS side by side on the same analytical system. In the past, such a study would have been challenging because chromatographic hardware was typically developed for specific flow rates. In turn, this would have necessitated complex hardware modifications to be able to achieve fair comparisons on a single analytical system. The recent commercialization of the Vanquish Neo LC system now allows such investigations because it can operate at flow rates ranging from 1 nL/min to 100 µL/min without any hardware changes. Therefore, the main purpose of the current study was to perform such a comparison and collect empirical data to provide guidance to the scientific community in selecting the most suitable flow rate systems for their specific applications.

## MATERIALS AND METHODS

### Sample Preparation

Human epithelial cervix carcinoma HeLa cells (ATCC, CCL-2) were cultured in Dulbecco’s Modified Eagle Medium (DMEM; Gibco, Invitrogen), supplemented with 10% fetal bovine serum, 100 U/mL penicillin (Invitrogen), 100 µg/mL streptomycin (Invitrogen), Cultures were incubated at 37 °C in a humidified atmosphere containing 5% CO_2_. Cells were harvested at approximately 80% confluence by washing twice with PBS and directly lysed on the culture plate using a buffer containing 8 M urea, 80 mM Tris-HCl (pH=7.6), 1×EDTA-free protease inhibitors (Complete Mini, Roche), and 1×phosphatase inhibitors (Sigma Aldrich). The plate was incubated on ice for 5 minutes before collecting the lysate by scraping. The lysate was centrifuged at 20,000×g at 4 °C for 30 minutes, and the supernatant was stored at −80 °C for further analysis. HeLa proteins were digested according to the SP3 protocol [15]. The obtained peptides were desalted using the HLB desalting cartridge. Peptides were quantified using Pierce Quantitative Fluorometric Peptide Assay, were dried using a SpeedVac, and stored at −20 °C.

Human body fluid specimens used in this study were obtained following informed consent and in accordance with the appropriate ethics approval process of the Technical University of Munich. Blood plasma (1 mL) was collected from a healthy donor and centrifuged at 4000 g for 10 minutes at 4 °C to obtain the supernatant. A 50 µL aliquot of the supernatant was diluted fivefold with 8 M urea containing 80 mM Tris-HCl (pH=7.6) the sample was further diluted with five volumes of 40 mM Tris-HCl buffer. For reduction, 1 M Dithiothreitol (DTT) was added to achieve a final concentration of 10 mM, and the mixture was incubated at 37 °C for 60 minutes. Alkylation was performed by adding chloroacetamide (CAA) to a final concentration of 55 mM, followed by incubation at room temperature in the dark for 30 minutes. Proteins were digested using a two-step trypsin digestion protocol with a protease-to-protein ratio of 1:100 (w/w) for each step. Desalting was carried out using the C18 StageTip protocol. Cerebrospinal fluid (CSF) samples were obtained from 10 individuals without neurological abnormalities, samples were pooled and stored at −80 °C until further use. Proteins were digested following the SP3 protocol described above for HeLa cells. Desalting was carried out using the C18 StageTip protocol. Peptides were quantified using Pierce Quantitative Fluorometric Peptide Assay.

### Phosphopeptide enrichment

In total 500 μg HeLa protein digest was separated on a 2.1 × 150 mm Waters XBridge BEH130 C18 3.5 µm column at a flow rate of 200 µl/min. Buffer A was 25 mM ammonium bicarbonate (pH =8.0), buffer C was 100% ultrapure water (ELGA), buffer D was 100% acetonitrile (ACN), buffer B was not used in this system. Separation was done with a linear gradient from 4% D to 32% D in 45 min, ramped to 80% D in 6 min, and kept there for 3 min before ramped back to 5% D in 2 min. 96 fractions were collected at 0.5 min intervals. Peptides were pooled in a step-wise fashion from 96-48 to 24-12 fractions according to the scheme above. Fractions were dried in a SpeedVac and stored at −80 °C until subsequent phosphopeptide enrichment. Phosphopeptides were enriched from each 12 fractions using Fe(III)-IMAC-NTA (Agilent Technologies) on the AssayMAP Bravo Platform (Agilent Technologies). IMAC cartridges were primed with 100 µl of 99.9% ACN/0.1% TFA and equilibrated with 50 µl loading buffer (80% ACN/0.1% TFA). Samples were reconstituted in 200 µl of loading buffer loaded onto cartridges, and washed with 50 µL loading buffer. Phosphopeptides were eluted with 40 µl of 1% ammonia, were quantified using NanoDrop^TM^ 2000 (Thermo Scientific) and were dried down and stored at −80 °C until subjected to LC-MS/MS analysis. 500 ng peptide loading was used per injection in both capLC and nLC-MS/MS systems.

### Kinobeads pull-downs

Kinobeads selectivity profiling of the multi-kinase inhibitor AT-9283 was conducted using a standard published protocol [16]. Briefly, K-562 (ATCC, CCL-243), COLO-205 (ATCC, CCL-222) and MV-4-11 (ATCC, CRL-9591) cells were cultured in RPMI 1640 medium (Biochrom GmbH) supplemented with 10% (v/v) FBS (Biochrom GmbH) and 1% (v/v) antibiotics. SK-N-BE(2) (ATCC, CRL-2271) cells were grown in DMEM/Ham’s F-12 (1:1) supplemented with 10% or 15% (v/v) FBS, respectively and 1% (v/v) antibiotics (Sigma). OVCAR-8 (RRID: CVCL_1629) cells were cultured in IMDM medium (Biochrom GmbH) supplemented with 10% (v/v) FBS. Cells were lysed in buffer containing 0.8% NP40, 50 mM Tris-HCl (pH=7.5), 5% glycerol, 1.5 mM MgCl_2_, 150 mM NaCl, 1 mM Na_3_VO_4_, 25 mM NaF, 1 mM DTT, protease inhibitors (SigmaFast), and phosphatase inhibitors (prepared in-house following Sigma-Aldrich’s cocktail 1,2 and 3 protocols). A pooled lysate (2.5 mg protein) from all five cell lines was pre-incubated with AT-9283 at increasing concentrations (DMSO vehicle, 3 nM, 10 nM, 30 nM, 100 nM, 300 nM, 1 µM, 3 µM, 30 µM) for 45 minutes at 4°C in an end-over-end shaker. Kinobeads (18 µL settled volume) were added to the lysate-compound mixture and incubated for 30 minutes at 4 °C with end-over-end agitation. After washing, bead-bound proteins were reduced with 50 mM DTT in 8 M Urea, 40 mM Tris-HCl (pH=7.4) for 30 minutes at room temperature, alkylated with 55 mM CAA, and digested with trypsin. Peptides were desalted using SepPak C18 µElution plates (Waters) and dried in a SpeedVac prior to LC-MS/MS analysis. 500 ng peptide loading was used per injection in both capLC and nLC-MS/MS systems.

### LC-MS/MS analysis

All LC-MS/MS analyses were performed on a Vanquish Neo LC system coupled to a Q Exactive HF-X mass spectrometer (Thermo Fisher Scientific) operating in positive polarity mode. Reverse-phase chromatography was performed using 150 mm long columns of varying i.d. (75, 150, 300, 1,000 µm) packed with identical material (Acclaim PepMap Neo C18, 2 μm particle size, 100 Å pore size, Thermo Fisher Scientific). Solvent A was 0.1% FA/3% DMSO (v/v) in water; solvent B was 0.1% FA/3% DMSO (v/v) in 100% ACN. DMSO was increased to 5% for flow rates <1.5 μL/min. The column temperature was maintained at 55 °C unless otherwise stated. Between runs, columns were equilibrated with three column volumes of 1% solvent A. A 25 μL sample loop was used for all direct injection setups and 10 μL volume was injected. Data-dependent acquisition (DDA) was employed using a full MS-ddMS2 method (S-lens RF level 40, 360-1300 m/z MS1 scan range, full MS AGC target 3E6, max injection time (IT) of 50 ms, for MS2 scans: 1.3 m/z isolation width, 100 m/z fixed first mass, max IT 22 ms, higher-energy collisional dissociation (HCD) fragmentation with a normalized collision energy (NCE) of 28, charge states from +2 to +5, peptide match set to preferred and isotope exclusion on. MS1 and MS2 spectra were acquired in profile and centroid mode, respectively. Further set-up specific details are provided below.

*µLC - 50 μL/min:* samples were analyzed using an Acclaim PepMap Neo C18 column (1 mm i.d. × 150 mm, P/N 164711) with a 54-min gradient (1-3.3% B in 1 min, 3.3-20% B in 45.1 min, 20-28% B in 6 min, 28-90% B in 0.9 min and wash at 90% B for 2 min) at 50 μL/min. Sample loading, equilibration and wash was done at 100 μL/min. 50 μm i.d. nanoViper capillaries connected the LC system to the Ion Max API source (HESI-II probe, depth set to A line). MS settings: spray voltage 4.0 kV, capillary temperature 320 °C, vaporizer temperature 200 °C, sheath/aux/sweep gas flow rates 32/5/0, full MS resolution 60,000 (at m/z 200), MS2 settings: resolution 15,000, intensity threshold 9E4, AGC target 1E5, Top12 method (28 Hz), dynamic exclusion 25 s.

*µLC - 10 μL/min:* samples were analyzed using an Acclaim PepMap Neo C18 column (300 μm i.d. × 150 mm, P/N 164537) with a 54-min gradient (same as 50 μL/min method above) but at 10 μL/min flow rate. Sample loading, equilibration and wash was performed at 15 μL/min. NanoViper capillary connections were same as for 50 μL/min setup. MS settings were identical to the 50 μL/min setup, except: spray voltage 2.5 kV, capillary temperature 320 °C, vaporizer temperature 72 °C, sheath/aux/sweep gas flow rates 8/2/0, Top18 method (28 Hz), dynamic exclusion set to 30 s.

*capLC - 5 μL/min:* Identical to the 10 μL/min setup, except: 5 μL/min flow rate during the gradient, spray voltage 2.3 kV, vaporizer temperature 63 °C, sheath/aux/sweep gas flow rates 8/1/0.

*capLC - 1.5 μL/min:* samples were analyzed using an Acclaim PepMap Neo C18 column (150 μm i.d. × 150 mm, P/N DNV150150PN) with a 50-min gradient (1-3.3% B in 1.5 min, 3.3-12% B in 26.1 min, 12-20% B in 15.5 min, 20-28% B in 4.5 min, 28-90% B in 0.8 min, and wash at 90% B for 1.6 min) at 1.5 μL/min. Sample loading, equilibration and washing was done at 3 μL/min. 20 μm i.d. nanoViper capillaries connected the pump, valves, and column. The column outlet was directly interfaced with the Nanospray Flex source (30 μm i.d. steel emitter) using a nanoViper to open silica capillary tube adapter (P/N 00109-02-00055). MS settings: spray voltage 3 kV, capillary temperature 275 °C, full MS resolution 60,000 (at m/z 200). MS2 settings: resolution 15,000, intensity threshold 9E4, AGC target 1E5, Top24 method (28 Hz), dynamic exclusion at 40 s. For the measurement of phosphopeptides, MS2 settings: max IT was changed to 50 ms and for analyzing kino-beads pulldown experiments Top12 method was used instead.

*nLC - 0.3 μL/min direct-injection:* samples were analyzed using an Acclaim PepMap Neo C18 DNV column (75 μm i.d. × 150 mm, P/N DNV75150PN) with a 45-min gradient (1-3% B in 3.3 min, 3-10% B in 22 min, 10-20% B in 14.7 min, 20-35% B in 3.3 min, 35-90% B in 0.5 min and wash at 90% B for 1.2 min) at 0.3 μL/min. Sample loading, equilibration and washing was done at 1.0 μL/min. 20 μm i.d. nanoViper capillaries connected the pump, valves, and column. The column outlet was interfaced with the Nanospray Flex Ion source (30 μm i.d. steel emitter) using a nanoViper to open silica capillary tube adapter (P/N 00109-02-00055). MS settings: spray voltage 2.1 kV, capillary temperature 275 °C, full MS resolution 120,000 (m/z 200). MS2 settings: resolution 15,000, intensity threshold 9E4, AGC target 1E5, Top24 method (28 Hz), dynamic exclusion 40 s. For the measurement of phosphopeptides, MS2 settings: max IT was changed to 50 ms and for analyzing kino-beads pulldown experiments Top12 method was utilized instead.

*nLC - 0.3 μL/min trap-and-elute:* samples were analyzed using an Acclaim PepMap Neo C18 SNV column (75 μm i.d. × 150 mm, P/N SNV75150PN) with a 56-min gradient (1-3% B in 4 min, 3-10% B in 23 min, 10-20% B in 22 min, 20-35% B in 5.9 min, 35-90% B in 0.5 min and wash at 90% B for 0.8 min) at 0.3 μL/min without heating. Sample loading was done on a trap cartridge (300 um i.d. × 5 mm, P/N 174500) at 50 μL/min in back-flush mode, column equilibration and washing was done at 0.5 μL/min. 20 μm i.d. nanoViper capillaries connected the pump, valves, and column inlet, the column outlet (open capillary end) was directly interfaced with the Nanospray Flex source (30 μm i.d. steel emitter) via an adaptor (P/N 13040420). MS parameters were identical to the direct-injection method above, except that dynamic exclusion was set to 30 s.

*Multi-Nozzle Setups:* for the 5 and 1.5 μL/min capLC setups, a MnESI source (Newomics) was used, employing the M3 8-nozzle 10 μm i.d. emitter. The MS was operated as follows: spray voltage 4.2 kV for 5 ul/min and 4.3kV for 1.5 ul/min; capillary temperature 320 °C; vaporizer temperature 30 °C. The flow rates of sheath gas, aux gas and sweep gas were set to 1, 0, and 0, respectively. The LC gradient for 5 μL/min multi-nozzle setup was reduced to a 52 min (3.3-20% B in 44 min, 20-28% B in 6 min, 28-90% B in 2 min) to accommodate slower sample loading, equilibration and washing speed at 10 μL/min, necessary for continuous spray stability. All other LC and MS parameters were identical to the respective flow rate setups above.

### Raw MS data processing and analysis

Raw data files were processed using MaxQuant (v1.6.2.10) [17] and searched against a human FASTA database (UniProtKB release 07.2019, UP000005640_74349) using default settings. Briefly, trypsin was set as the enzyme with up to two missed cleavages. Cysteine carbamidomethylation was configured as a fixed modification, while N-terminal acetylation and methionine oxidation were specified as variable modifications. The false discovery rate (FDR) was maintained at 1% at the site, peptide-spectrum match (PSM), and protein levels. Chromatographic full width at half-maximum (FWHM) information was extracted from the MaxQuant output file “allPeptides.txt”. Andromeda scores were taken from the “evidence.txt” after excluding all reverse sequences and potential contaminants. Peptide intensity boost comparisons were evaluated from “peptides.txt” using the Benjamini-Hochberg multiple testing correction procedure [18] set to 0.05 FDR. For analyzing kinobeads pulldown data, the CurveCurator pipeline [19] was employed, permitting up to six missing values, using a fold change minimum of 0.5 and an alpha value cutoff of 0.01. pEC_50_ values (=-log_10_EC_50_) were obtained from CurveCurator analysis, apparent dissociation constants (Kd^app^; often expressed as pKd^app^=-log_10_Kd^app^) were subsequently calculated by multiplying EC_50_ values by correction coefficients, defined as the ratio of pulldown of pulldowns experiment intensity values to DMSO control. Data analysis and visualization were performed using in-house developed R and Python scripts, along with Skyline (v22.2) [20], Biorender and Inkscape.

## RESULTS AND DISCUSSION

### Study design for the side-by-side performance comparison of proteome analysis by µLC-, capLC- and nLC-MS/MS

The overall design of this comparison is summarized in **Figure 1A**. The key principles driving this design were to perform all experiments using i) the same samples, ii) the same HPLC, iii) the same column material and length, iv) the same mass spectrometer, v) the same sample throughput and vi) the same analysis software.

**Figure 1.**
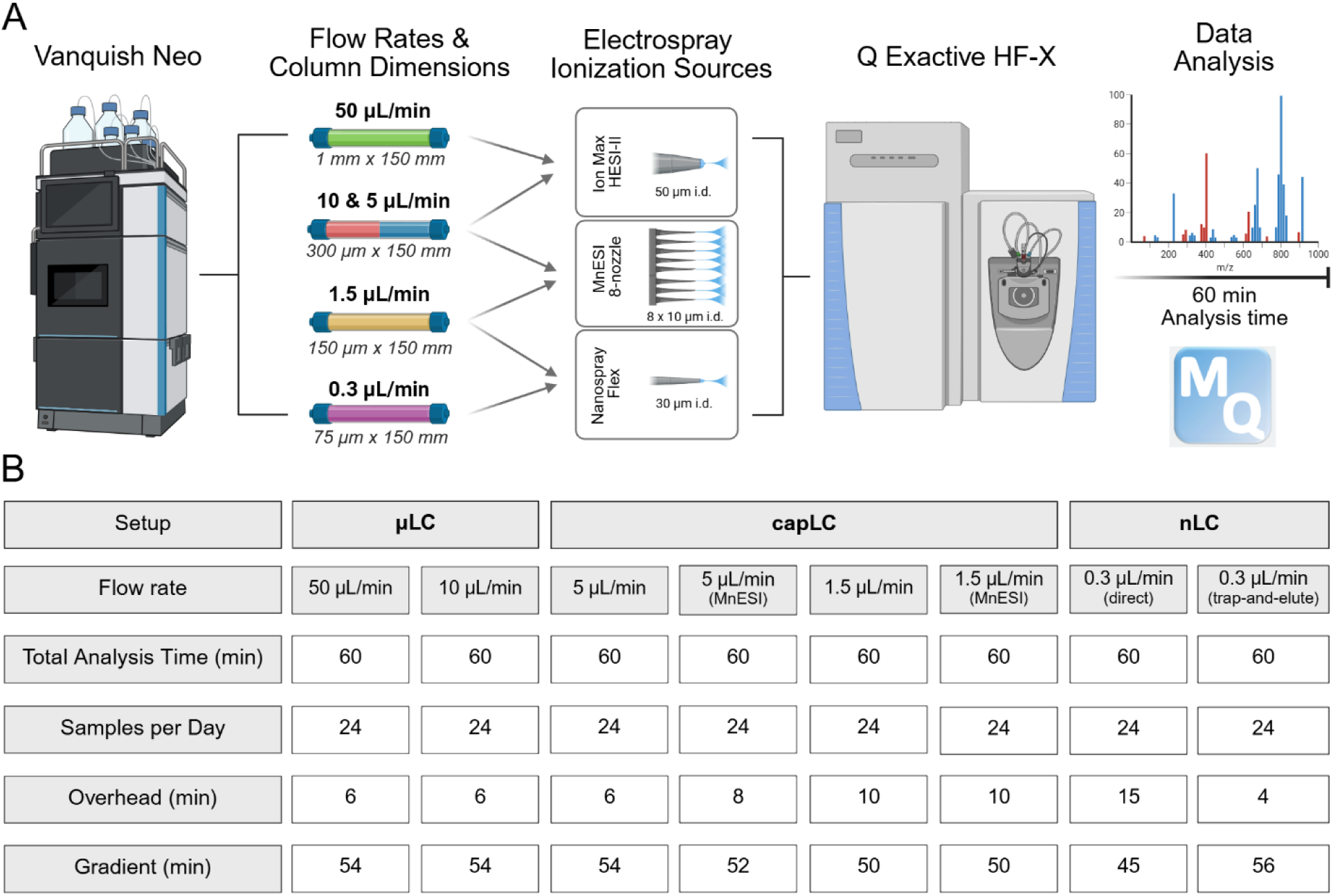
(A) Study design for the comparative performance evaluation of µLC-, capLC- and nLC-MS/MS systems for proteome analysis. (B) Tabular summary of key LC parameters for each setup.

The Vanquish Neo UHPLC system accommodated all five flow rates tested in this study and all separations were performed using commercially available Acclaim PepMap Neo C18 columns as follows: µLC (50 μL/min: 1 mm i.d. × 150 mm; 10 μL/min: 300 μm i.d. × 150 mm), capLC (5 μL/min: 300 μm i.d. × 150 mm; 1.5 μL/min: 150 μm i.d. × 150 mm) and nLC (0.3 μL/min: 75 μm i.d. × 150 mm). All analytical columns had the same length, featured identical particle size and chemistry (2 μm/100 Å, C18) and only differed by internal diameter. Depending on the setup, coupling of the Vanquish Neo with the Q Exactive HF-X mass spectrometer was achieved by using either the Ion Max API Ion source with a HESI-II probe (for 50, 10 and 5 μL/min flow rates) or the Nanospray Flex Ion source (for 1.5 and 0.3 μL/min flow rates). An add-on to the study was to employ a commercial multi-nozzle electrospray emitter for the 1.5 and 5 μL/min flow rates.

To assess sensitivity of each setup, serial dilutions of a HeLa cell line digest (5 ng - 10 μg on column) were analyzed in triplicate. Instead of fixing the analytical gradient to a uniform length, all comparisons were made using the increasingly popular “samples per day” (SPD) approach. This concept proved to be practical particularly when planning larger-scale projects. To allow for comprehensive sampling, the injection-to-injection time was fixed to 60 minutes (24 SPD) for each flow rate throughout this study. The LC systems were operated in direct sample injection mode, except for nLC, where both direct and trap-and-elute injection modes were compared. A consequence of using the SPD approach was that the analytical gradient times and overhead times (sample loading and column equilibration) were not the same for all setups (**Figure 1B**) because of the longer sample (10 μL volume) loading and equilibration times required for the direct injection gradient flow rates of 0.3 and 1.5 μL/min. This effect did not apply to the trap-and-elute setup because sample loading on the trap column was performed at a much higher flow rate of 50 μL/min. LC and MS parameters were individually optimized for each flow rate and gradient shapes were tailored to make separations as uniform as possible.

### Comparison of the sensitivity of proteome analysis by µLC-, capLC- and nLC-MS/MS

The sensitivity of each setup was assessed by analyzing serial dilutions of HeLa cell line digests in triplicate, initially using the number of identified peptides and proteins as a metric (**Figure 2A,B**). The observed general trends align well with expectation such that lower flow rates (≤1.5 μL/min) outperform higher flow rates for peptide loadings below 1 μg where electrospray ionization (ESI) efficiency drives sensitivity. Conversely, higher flow rate separations require higher sample loading to compensate for the loss of ESI efficiency. The observed increase in identified peptides and proteins with increasing loading amounts allowed us to determine an optimal range of peptide loading for each flow rate setup. The observed saturation serves as a valuable indicator for estimating the upper limit of the appropriate sample amount for each system. Identifying the optimum range is of practical value, as injecting too little results in lower IDs, while injecting too much can exceed column capacity, leading to deterioration of chromatographic peak shapes, fewer IDs and poorer quantification [22]. Based on the empirical data, 10-50 μL/min flow rates are best for sample loadings of ≥5 μg and 5 μL/min is effective for 1-5 μg. Interestingly, 1.5 and 0.3 (trap-and-elute) μL/min showed very similar performance in the 10 ng to 1 μg range with consistently higher numbers for the 1.5 μL/min capLC system starting from 200 ng loading. As a second metric for comparing the sensitivity of the different systems, we used the chromatographically integrated peptide precursor ion intensity measured by the mass spectrometer. To ensure fair comparison, we fixed sample loading to 200 ng ensuring that all systems were below saturation point. The 50 μL/min setup was used as a reference and peptides detected in this setup were used to calculate the intensity fold-changes of peptides between the different flow rates (FDR=0.05, Benjamini-Hochberg procedure). As shown in **Figure 2C**, reducing the flow rate from 50 to 10 μL/min resulted in a median 1.7-fold increase in peptide intensity. Further reduction to 5 μL/min yielded an almost 4-fold increase. A major change occurred at a flow rate of 1.5 μL/min which achieved a 14-fold increase in intensity and a further improvement to 20 to 23-fold was reached using nLC operated at 0.3 μL/min. At higher sample loadings, the intensity boosts observed at lower flow rates strongly diminished. This is expected because at lower flow rates, the ESI process saturates while higher flow rates benefit from higher sample loading to offset the poorer ESI efficiency (**Figure S.1**).

**Figure 2.**
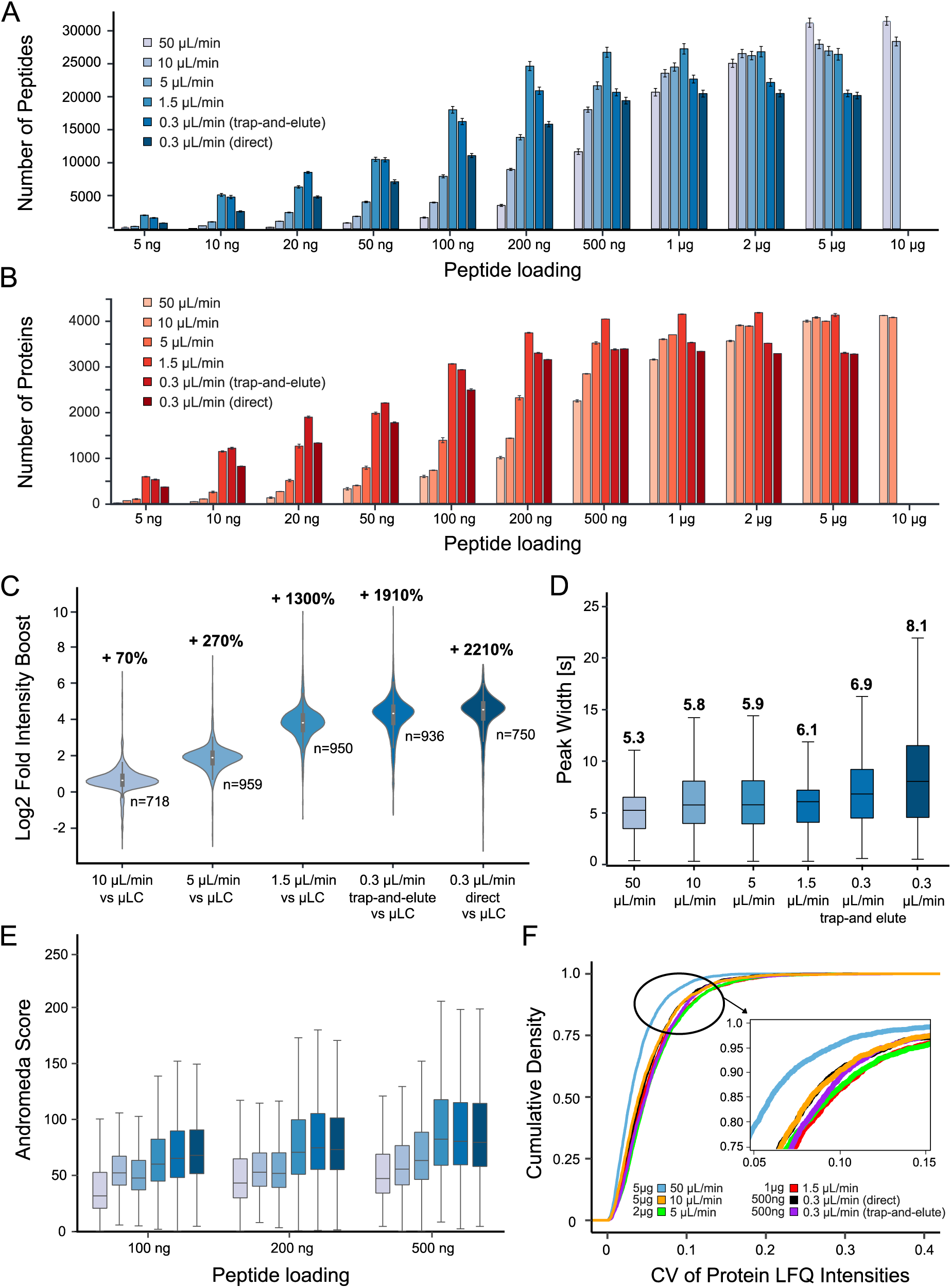
Comparison of all LC-MS/MS setups based on serial dilution experiments (n=3 measurements for each sample loading). (A) Bar plot showing the average number of identified peptides for each sample loading. (B) Same as (A) but for protein groups (red). (C) Violin plots showing the distribution of the relative boost of peptide intensities (based on extracted ion chromatograms) for all setups at 200 ng peptide loading compared to the reference µLC setup operating at a flow rate of 50 μL/min (n=number of peptides used for comparison; color code as in (A)). (D) Box blots showing the distributions, medians and interquartile ranges of chromatographic peak widths at half maximal signal (FWHM) of all identified peptides at optimal column loadings. Color code as in (A). (E) Box blots showing the distributions, medians and interquartile ranges of MaxQuant Andromeda scores of all setups at three levels of peptide loading. Color code as in (A). (F) Cumulative density plot illustrating the quantitative repeatability (precision) of all setups measured by the coefficient of variation (CV) of quantified proteins when using optimal peptide loadings.

The observation that the 1.5 μL/min capLC setup was on par or even outperformed the 0.3 μL/min nLC trap-elute setup in terms of peptide and protein identifications was surprising given that the nLC system exhibited higher sensitivity (∼42% higher median peptide intensity) and benefitted from four more minutes of analytical gradient time (8%) because of the shorter overhead time for sample loading and system equilibration. Still, capLC prevailed, most probably because of its superior chromatography, evidenced by substantially sharper peaks (median FWHM of 6.1 vs 6.9 s; 13%) as well as a much narrower distribution of LC peak widths (**Figure 2D**) both of which increase chromatographic peak capacity. The improved separation also comes with the effect of a higher peptide concentration that improves ESI response particularly of low abundance peptides, in turn leading to higher signal intensities, thereby increasing the probability of triggering a data-dependent scan and doing so closer to the apex of the LC peak [22]. Higher peptide signal also results in more precursor ions for fragmentation within a given time frame leading to enhanced quality of MS2 spectra and, ultimately, more peptide and protein identifications. This interpretation is supported by the very similar Andromeda score distributions of identified peptides of the 1.5 μL/min capLC and 0.3 μL/min nLC systems starting from 200 ng loading (**Figure 2E, Figure S.1B)**

Robust peptide and protein quantification in proteomic experiments is at least if not more important than covering a lot of proteins. The 50 μL/min system showed the best overall quantitative precision with a coefficient of variation (CV) of <10% for >95% of all quantified proteins. Still, when adjusting peptide loading to an appropriate level, all evaluated chromatographic setups achieved excellent quantitative precision (<20% CV for >95% of all quantified proteins; **Figure 2G**).

### Performance evaluation of a multi-nozzle ESI source

To evaluate if the performance of the capLC systems (1.5 and 5 μL/min) could be further improved, we tested a commercial multi-nozzle ESI source (MnESI, Newomics) which splits the LC flow into eight streams, effectively reducing the flow delivered to each emitter nozzle by 8-fold (M3 Emitter). The underlying concept is to enhance ESI efficiency by reducing flow rate post-column while maintaining full chromatographic performance. Kreimer et al. recently reported results using such a multi-nozzle emitter in conjunction with a 300 μm i.d. × 50 mm C8 column running at 9.5 μL/min and a dual trap setup to analyze plasma and cell digest samples by data independent acquisition (DIA) on a timsTOF mass spectrometer at a rate of 15 min/sample. They identified 400 proteins in plasma and 4,000 proteins in cell digests. However, no comparison to a reference system without a multi-nozzle emitter was provided [23].

Here, we again used serial dilutions of Hela cell digests to compare the capLC setups with and without a multi-nozzle emitter side by side. The 5 μL/min setup showed a modest benefit in the number of peptide (up to 16%) and protein (up to 18%) identifications (**Figure 3A,B**) even though the available gradient time was two minutes shorter than on the reference system. This benefit was, however, not observed at sample loadings below 200 ng and diminished at high sample loads. For the 1.5 μL/min setup, the MnESI source performed substantially worse in terms of peptide and protein identifications than the reference NanoFlex source (**Figure 3C,D**) which was also reflected by lower median peptide intensities (**Figure 3E**). Chromatographic performance of MnESI and reference systems were near identical as expected (**Figure 3F)** and also no strong differences were detected when comparing quantitative precision (**Figure 3G**). Therefore, the observed differences above can be attributed to the ionization source itself. At a flow rate of 5 μL/min, the MnESI reduces flow to 0.625 μL/min per nozzle, possibly enhancing ESI efficiency. An additional or alternative contributing factor is that the MnESI’s emitter nozzles have a 10 μm i.d. tip opening, while the one of the HESI needle is 50 μm. The smaller emitter i.d. should work in tandem with the reduced flow rate to promote smaller droplet formation, thereby improving ESI efficiency. In contrast, for the 1.5 μL/min setup, transitioning from a NanoFlex 30 μm i.d. steel emitter to 10 μm i.d. multi-nozzle emitters proved ineffective. At a flow rate of 1.5 μL/min, the MnESI reduces the flow per nozzle to 0.188 μL/min. It is possible that the 10 μm i.d. emitter architecture is no longer efficient for a flow rate this low. Instead, a smaller internal diameter, such as 5 μm, may be required to produce appropriately sized droplets for these flow conditions but these were not commercially available.

**Figure 3.**
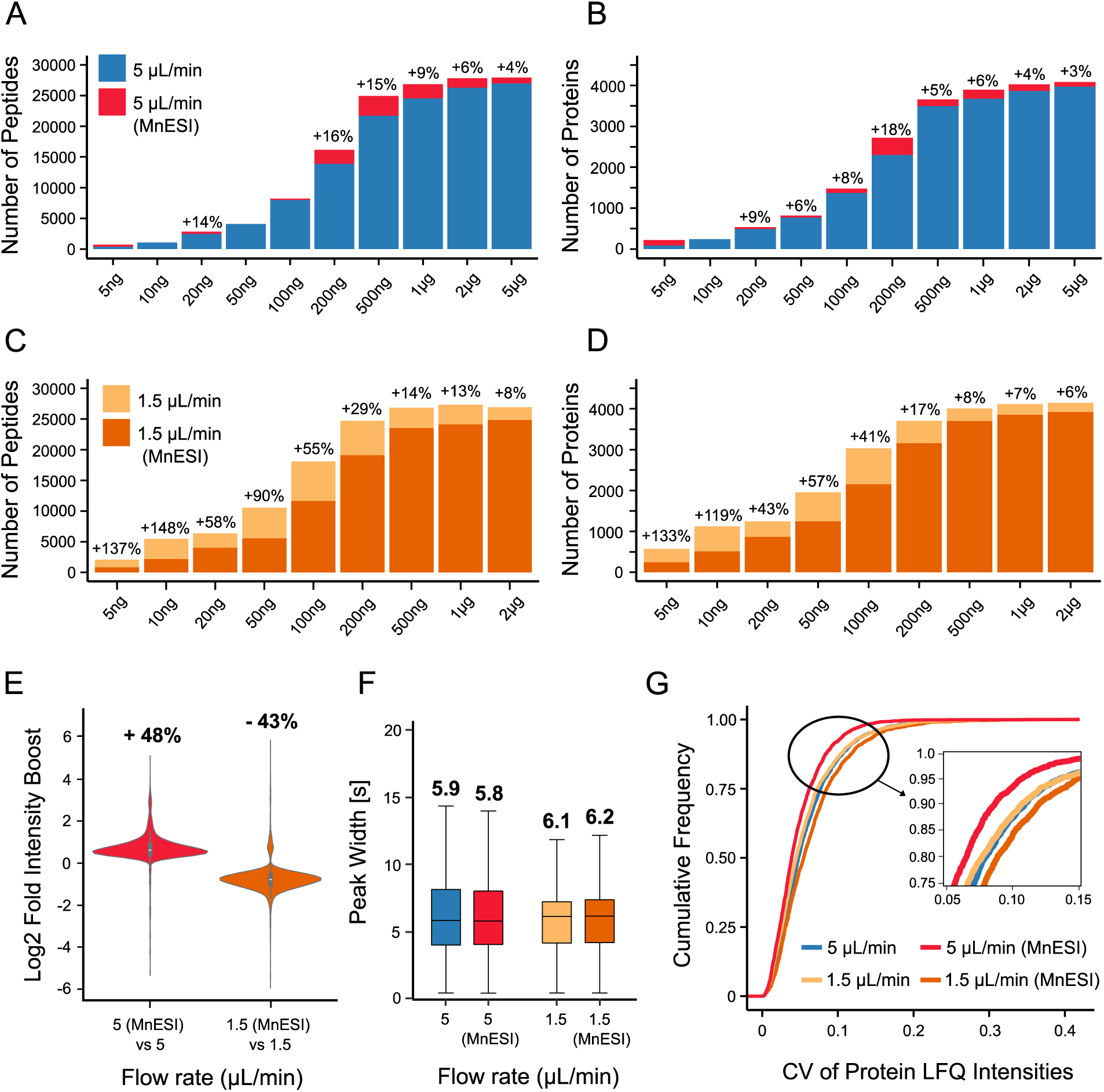
Benchmarking results of multi-nozzle electrospray ionization (MnESI) setups (A) Stacked bar plot showing the average number of identified peptides for each sample loading for 5 μL/min flow rates with and without using a MnESI source. (B) Same as (A) but for protein groups. (C) Same as (A) but for 1.5 μL/min (D) Same as (B) but for 1.5 μL/min. (E) Violin plots showing the distribution of the relative boost of peptide intensities (based on extracted ion chromatograms) of MnESI compared to the reference setups (optimal peptide loadings of 2 μg for the 5 μL/min and 1 μg for the 1.5 μL/min setups were used) (F) Box blots showing the distributions, medians and interquartile ranges of chromatographic peak widths at half maximal signal (FWHM) of identified peptides at optimal loading. (G) Cumulative density plot illustrating the quantitative repeatability (precision) of the capLC setups with and without multi-nozzle emitters at optimal peptide loading.

### Robustness and reproducibility of the 1.5 μL/min capLC-MS/MS setup

Given the overall very good performance of the 1.5 μL/min capLC setup, and the fact that this flow rate has been underexplored in the proteomics field, all further experiments were performed using this configuration. To access its technical robustness and reproducibility, we conducted an experiment consisting of 100 consecutive injections, divided into four identical cycles of 25 injections each (**Figure 4A**), including different sample types and spanning 4.5 days of measurement. Each cycle included 11 replicates of the same HeLa cell line digest, four replicates of the same cerebrospinal fluid (CSF) digest, and 10 replicates of the same human plasma digest. For each sample, 1 μg peptide loading was used and 300 fmol of a synthetic peptide retention time standard mix (PROCAL) [24] was spiked into each sample to monitor retention time stability. Blank runs were included between each sample type to evaluate sample carry-over. Analysis of PROCAL peptide retention times revealed peptide-specific CVs of 0.2-1.8% (average 0.7%), demonstrating high chromatographic reproducibility (**Figure 4B**). Excellent separation robustness, in turn also resulted in high reproducibility of the number of peptide (CV<2%) and protein group (CV<3%) identifications (**Figure 4C,D**). Identifications achieved for plasma were comparable to state-of-the-art single shot, unfractionated sample analysis reported in the literature [25]. For CSF, the number of identified peptides and proteins was almost twice as high as reported in a recent ring trial study [26]. This may be attributable to the higher loading amount used here (1 ug vs 400 ng), the fact that we pooled CSF from 10 donors and the possibility that our CSF samples were not free of tissue material that could easily be introduced during lumbar CSF collection. Quantitative precision was also outstanding, with CVs of <20% for 96% of quantified proteins for Hela and 98% for CSF and plasma samples (**Figure 4E**). In addition, sample carry-over was very low (0.12% peptide intensity for HeLa, 0.11% for CSF, and 0.48% for Plasma) demonstrating that the 1.5 μL/min capLC system is a highly effective setup for analyzing these sample types when using optimal sample loading.

**Figure 4.**
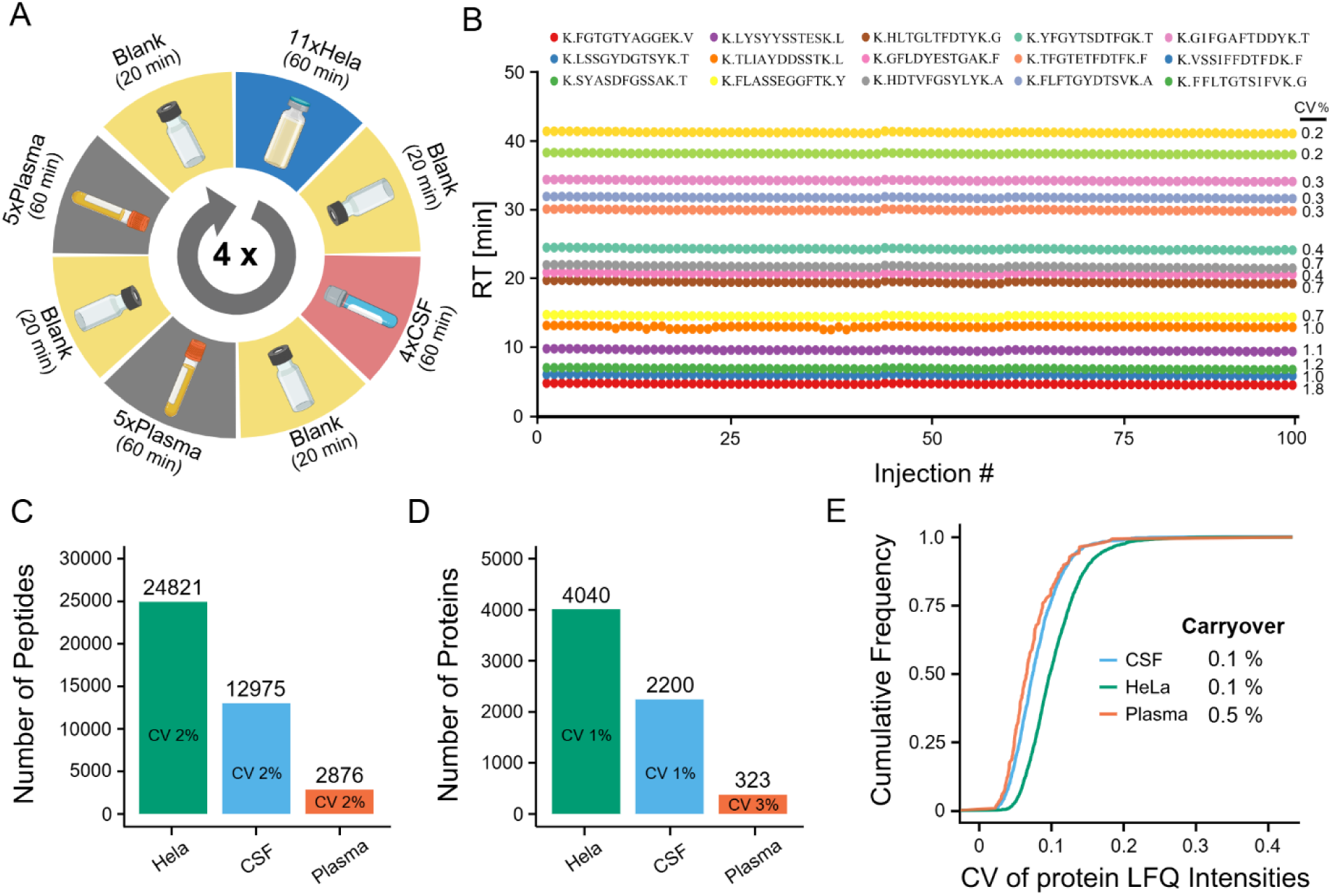
Robustness and reproducibility of the 1.5 μL/min capLC setup. (A) Scheme illustrating the design of the robustness and reproducibility test. (B) Retention time (RT) plots of 15 PROCAL peptides and their associated average retention time CVs across 100 consecutive sample injections. (C) Bar plot summarizing peptide identification stability across the cycle shown in (A). (D) Same as (C) but for protein groups. (E) Cumulative density plot illustrating the quantitative reproducibility (precision) as well as run-to-run average sample carryover of the different sample types.

### Example applications for capLC-MS/MS in proteomics

The above shows that the 1.5 μL/min capLC system can be used to analyze samples as different as plasma or cell lines. However, for plasma in particular, it can be argued that μLC is the more convenient choice because protein is easily and abundantly available (∼60 μg/μL) and the absolute sample loading required to achieve good protein coverage is far below the capacity of 1 mm i.d. columns, in turn ensuring high performance separations over long periods of time. At the other end of the spectrum, the proteome analysis of rare or single cell population will require nLC because of absolute sensitivity demands. An interesting middle ground is the analysis of sub-proteomes that can be enriched biochemically and two illustrative cases are exemplified below. In the first application, we purified phosphopeptides from 500 ug of HeLa cell line digest by immobilized metal affinity chromatography (IMAC; two workflow replicates). Three samples of 500 ng each were analyzed using either the capLC or direct injection nLC-MS/MS systems. Interestingly, the capLC setup identified ∼10,000 phosphopeptides, outperforming the nLC setup by 21% (11% at the level of phosphoproteins; **Figure 5A,B**). This difference is likely due to the five minutes shorter gradient time available on the direct injection nLC setup compared to the capLC system. As already mentioned above, the sharper chromatographic peaks of the capLC setup also contribute to this performance advantage (**Figure 5C**) as peptide intensities as well as Andromeda scores were comparable between the two setups (**Figure S.2A; Figure 5D**).

**Figure 5.**
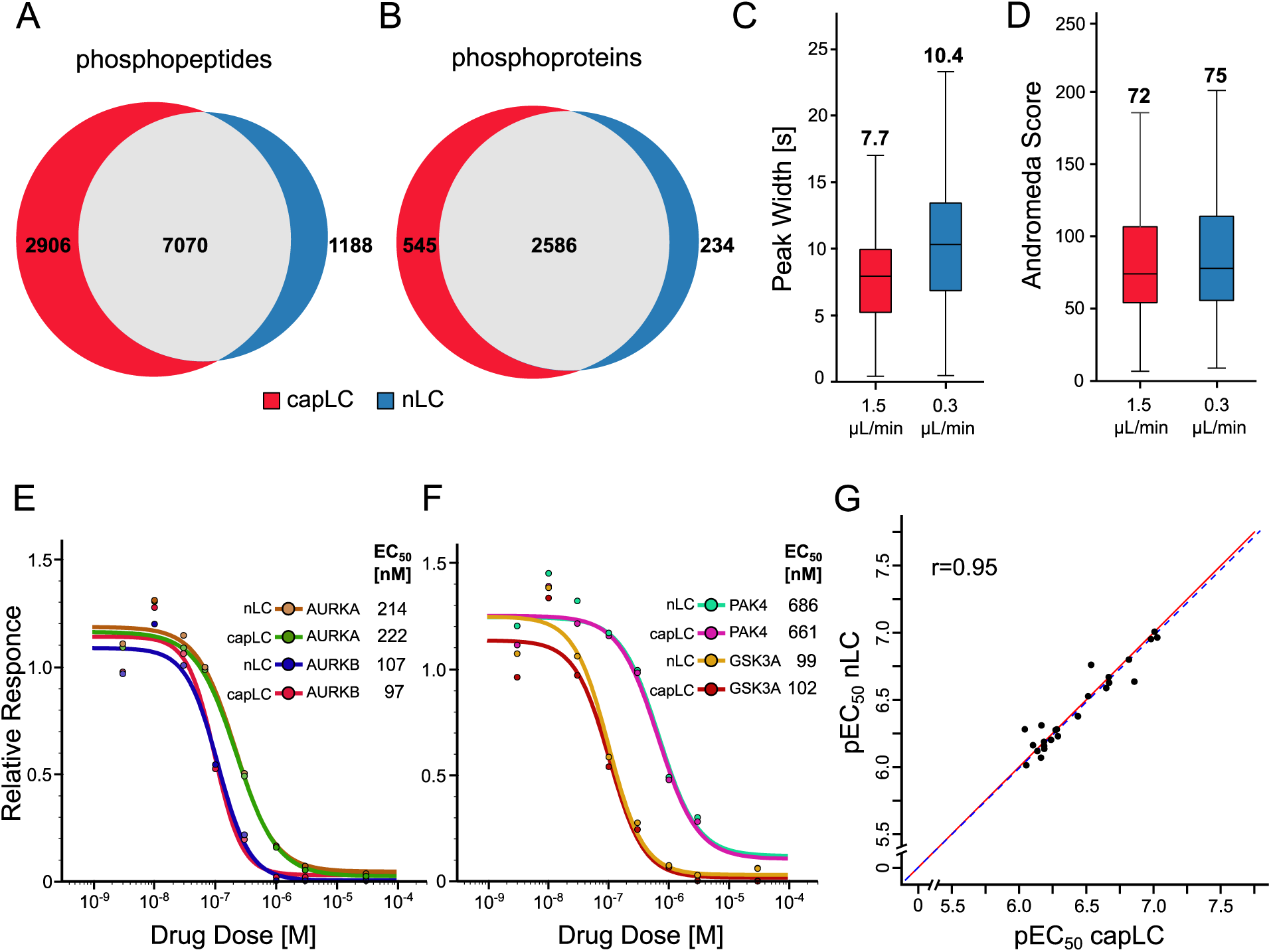
Comparative analysis of sub-proteomes on the 1.5 μL/min capLC and nLC systems (A) Venn diagram comparing the number of identified phosphopeptides. (B) Same as (A) but for phosphoproteins. (C) Box blots showing the distributions, medians and interquartile ranges of chromatographic peak widths at half maximal signal (FWHM) of identified phosphopeptides. (D) Box blots showing the distributions, medians and interquartile ranges of MaxQuant Andromeda scores of all phosphopeptides. (E,F) Dose-response curves for illustrating the interaction of the kinase inhibitor AT-9283 with a number of target proteins (EC_50_: effective concentration to reduce 50% of protein binding to Kinobeads). (G) Scatter plot correlating pEC_50_ values ((≥6; equivalent to EC_50_<1,000 nM) measured for drug-target interactions obtained by the capLC and nLC setups. r=Pearson correlation coefficient, fitted regression line in blue and x-y diagonal line in red.

In a second example, we compared the capLC to nLC-MS/MS setups for analyzing drug-protein interactions using the multi-kinase inhibitor AT-9283 and the Kinobeads approach [27]. The capLC and nLC systems identified 289 and 287 kinases respectively (**Figure S.2B**). CurveCurator [19] was used to analyze the dose-response data and to determine apparent interaction constants (K_d_^app^) for the compound and its target proteins. This analysis resulted in the identification of 58 and 54 targets using capLC-or nLC-MS/MS respectively (**Figure S.2C**). More importantly, overlaying the dose-response curves obtained by both analytical setups showed nearly identical results for the potent drug targets (e. g. AURKA, AURKB, GSK3A, and PAK4; **Figure 5E,F; Figure S.3**). Consequently, their determined half-maximal effective concentration (EC_50_) and K_d_^app^ values values were also very similar. This was also more generally true when correlating the pEC_50_ values (-log_10_EC_50;_ Pearson correlation coefficient r=0.95; **Figure 5G**) or pK_d_^app^ values (−log_10_K_d_^app^; r=0.88, **Figure S.2D**) obtained by capLC or nLC-MS/MS respectively for all commonly identified targets.

## CONCLUSIONS

In this study, we conducted a side-by-side performance evaluation of μLC, capLC, and nLC-MS/MS systems (300 nL/min to 50 uL/min) for use in proteomics. The data confirmed many previously reported results [6, 14] but also closed a gap in the literature by systematically evaluating all flow rates and column dimensions useful for standard proteomic applications and largely eliminating the influence of the particular sample preparation, mass spectrometer or data analysis software employed. A limitation of the current study is that it did not evaluate higher than 24 SPD throughput scenarios. However, we consciously chose to perform the evaluation using a 60 min turnaround time because the effective gradient times of between 45-56 min are long enough to ensure that all the data is of high qualitative, and, more importantly, quantitative quality. We also consciously chose a DDA over a DIA approach and refrained from employing performance enhancing data processing tools such as match-between-runs [28] or AI-based identification re-scoring approaches such as PROSIT [29] across different flow rate or sample loadings for the sake of being conservative and avoiding additional parameters that are unrelated to the chromatographic part of the overall analytical proteome analysis workflow. Still, the authors anticipate that the observations and learnings made for the 24 SPD data will translate to shorter gradients, as long as the employed mass spectrometer can keep up with the sharper LC peaks and higher number of co-eluting peptides at shorter gradients [30–32]. The commercial availability of HPLC systems that can accommodate a wide range of flow rates facilitates the choice of the right column and gradient for the right application especially for laboratories that cannot reserve dedicated LC-MS/MS systems for particular applications. We also anticipate that this study will provide useful practical guidance for scientists working in the field as to which setup to choose for a given application.

## ASSOCIATED CONTENT

### Supporting Information

The following files are available.

Supplementary Figures; it contains Supplementary figures that are referenced in the main text (file type is .pdf)

Supplementary Source Data; it contains source data used for creating Figures 2 A,B; 3 A,B,C,D; 4 B,C,D,E; 5G, and Supplementary Figure 2 D (file type is .xls)

## AUTHOR INFORMATION

### Author Contributions

B.K., R.Z. and C.P. conceived the study. G.T. and B.K. designed experiments. G.T., N.K., P.P performed experiments. G.T. analyzed the data. G.T. and B.K. wrote the manuscript.

## ACKNOWLEDGMENTS

This work was supported by the Elite Network of Bavaria (grant F.6-M5613.6.K-NW-2021-411/1/1) and the German Ministry for Science and Education (grants 03LW0243K and 16LW0243K). The authors are grateful to all members of the Kusterlab for technical assistance and fruitful discussions.

## ABBREVIATIONS

ACN: Acetonitrile
CAA: Chloroacetamide
capLC: Capillary Liquid Chromatography
CSF: Cerebrospinal Fluid
CV: Coefficient of Variation
DDA: Data-dependent Acquisition
DIA: Data-Independent Acquisition
DMSO: Dimethyl Sulfoxide
DTT: Dithiothreitol
EC50: Effective Concentration to Reduce 50% of Protein Binding to Kinobeads
FBS: Fetal Bovine Serum
fmol: Femtomole
FDR: False Discovery Rate
FWHM: Full Width at Half Maximum
HeLa: Human Cervical Carcinoma Cell Line
HLB: Hydrophilic-Lipophilic Balance
HPLC: High-Performance Liquid Chromatography
i.d.: Internal Diameter
Kdapp: Apparent Dissociation Constant
LC: Liquid Chromatography
LC-MS/MS: Liquid Chromatography-Tandem Mass Spectrometry
µg: Microgram
min: Minute
µL: Microliter
µLC: Microflow Liquid Chromatography
mm: Milimeter
mM: Milimolar
MS: Mass Spectrometry
ng: Nanogram
nL: Nanoliter
nLC: Nanoflow Liquid Chromatography
nM: Nanomolar
PSM: Peptide-spectrum Match
RT: Retention Time
SPD: Samples per Day
SWATH-MS: Sequential Window Acquisition of All Theoretical Mass Spectra
TFA: Trifluoroacetic Acid
Å: Angstrom

