## Supplementary Figures for "High-performance proteomics at any chromatographic flow rate"

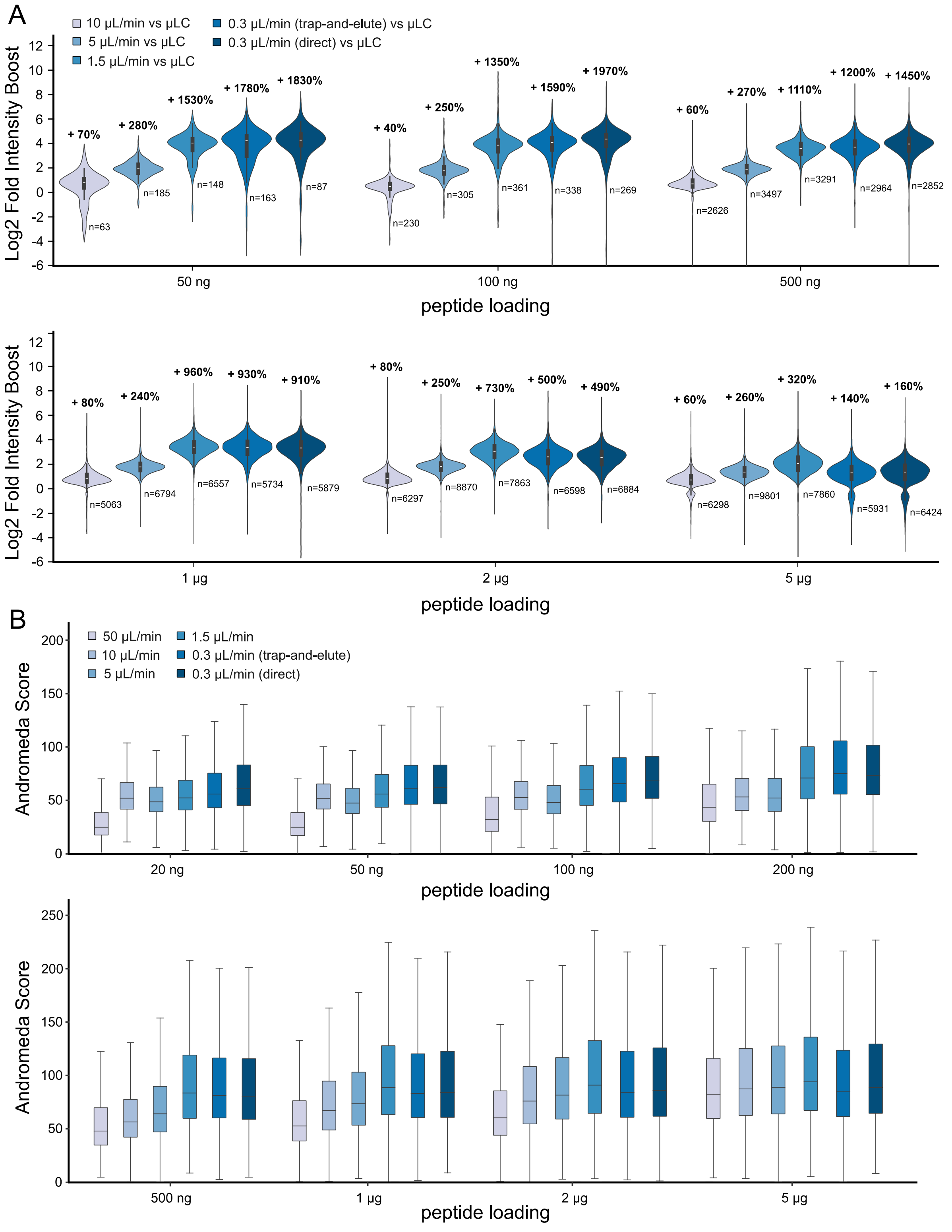


**Supplementary Figure 1.** (A) Violin plots showing the relative boost of peptide intensities (based on extracted ion chromatograms) and their intensity distributions for all setups at different peptide loadings compared to the reference µLC setup operating at a flow rate of 50 µL/min. "n" denotes the number of shared peptides used for analysis. (B) Box blots showing the distributions, medians and interquartile ranges of MaxQuant Andromeda scores of all setups at different peptide loadings.

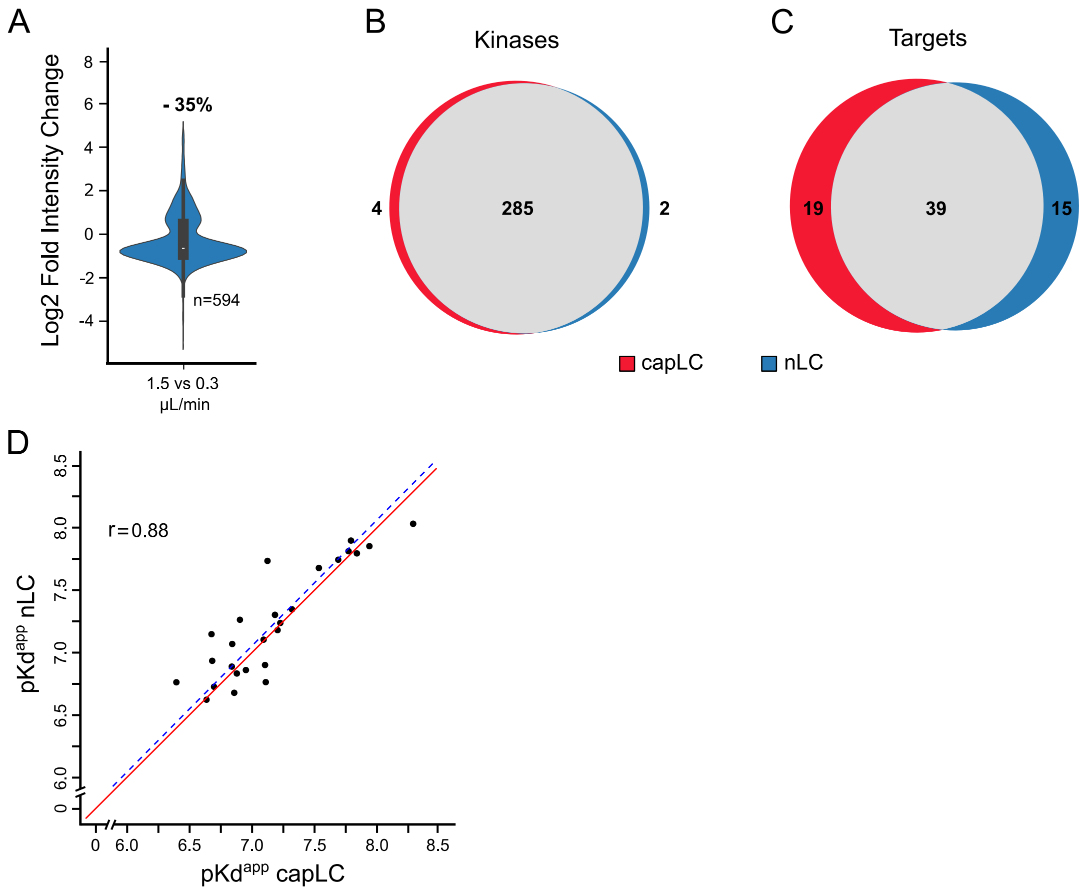


**Supplementary Figure 2.** (A) Violin plot showing the relative boost of peptide intensities (based on extracted ion chromatograms) and their intensity distribution for the capLC (1.5 µL/min) vs. nLC (0.3 µL/min) setup ("n" denotes the number of shared peptides used for analysis). (B) Venn diagram comparing the number of detected kinases in Kinobeads pulldown experiments using the kinase inhibitor AT-9283. (C) Same as (B) but counting only the target proteins of AT-9283. (D) Scatter plot correlating $pK_d^{app}$ values calculated from dose-response curves for drug-target interactions obtained by the capLC and nLC setups, r denotes Person correlation coefficient, fitted regression line in blue and x-y diagonal line in red.

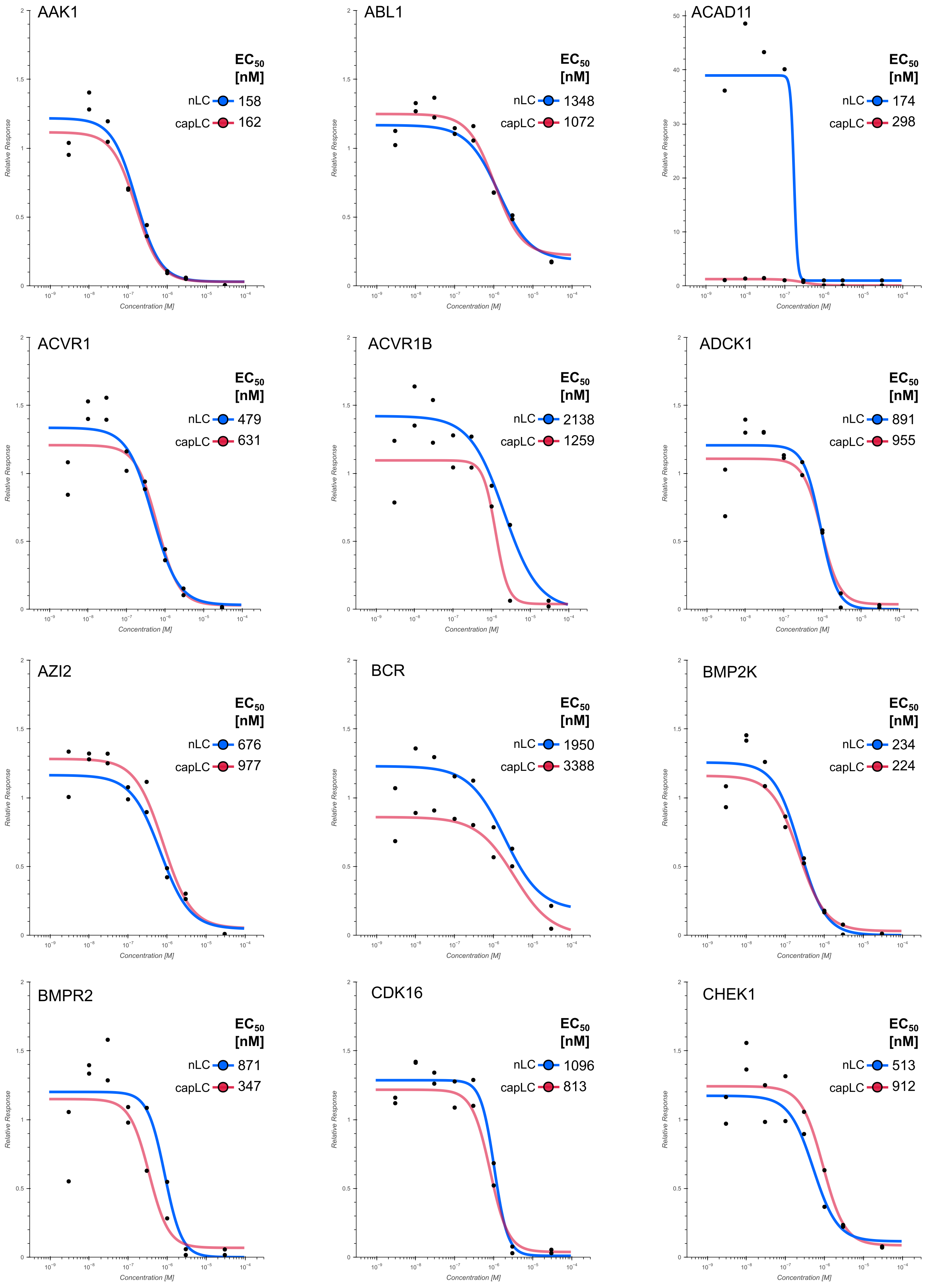
AAK1
EC50 [nM]
nLC 158
capLC 162
ABL1
EC50 [nM]
nLC 1348
capLC 1072
ACAD11
EC50 [nM]
nLC 174
capLC 298
ACVR1
EC50 [nM]
nLC 479
capLC 631
ACVR1B
EC50 [nM]
nLC 2138
capLC 1259
ADCK1
EC50 [nM]
nLC 891
capLC 955
AZI2
EC50 [nM]
nLC 676
capLC 977
BCR
EC50 [nM]
nLC 1950
capLC 3388
BMP2K
EC50 [nM]
nLC 234
capLC 224
BMPR2
EC50 [nM]
nLC 871
capLC 347
CDK16
EC50 [nM]
nLC 1096
capLC 813
CHEK1
EC50 [nM]
nLC 513
capLC 912
Relative Response
Concentration [M]

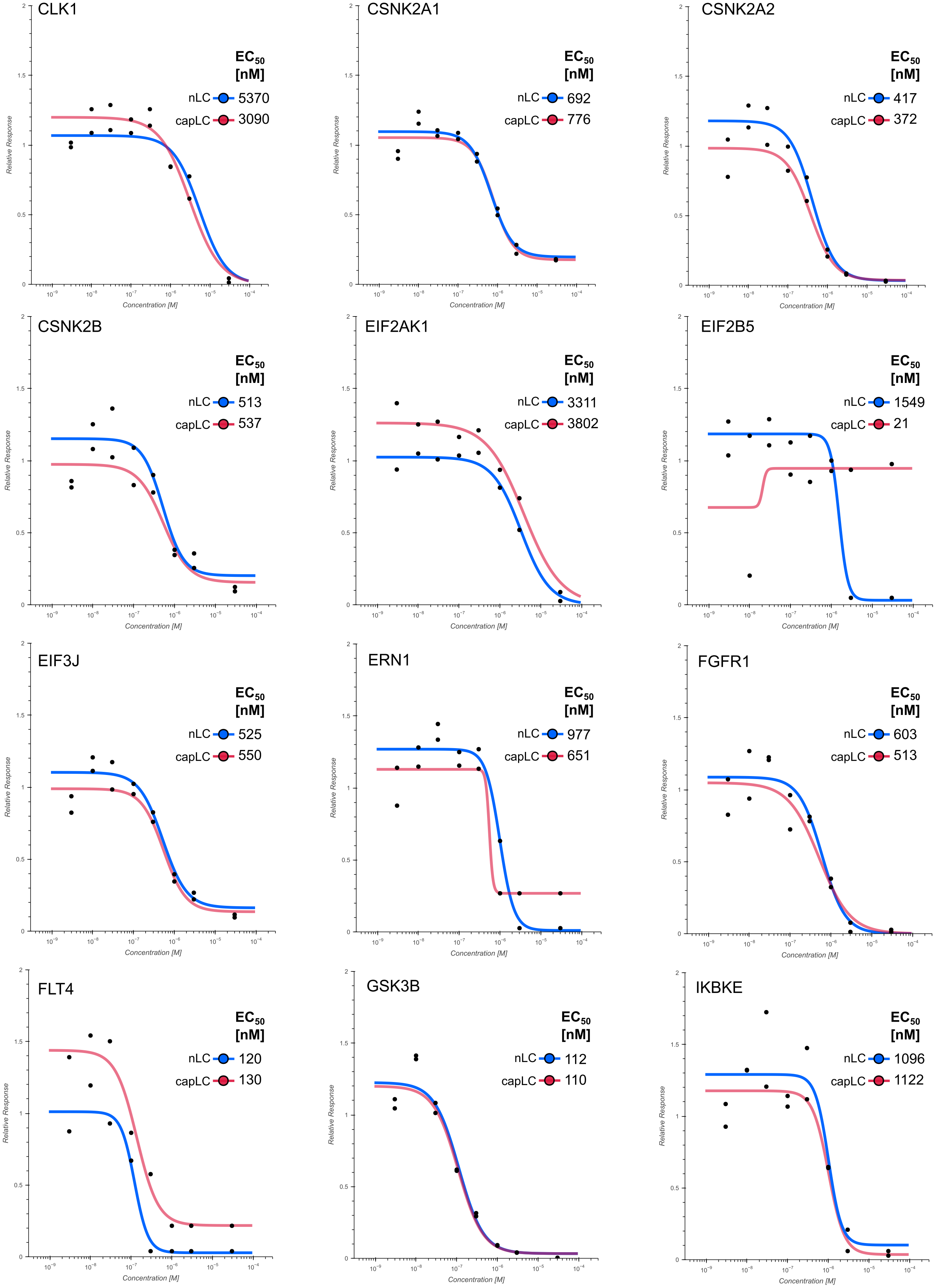
CLK1
EC50 [nM]
nLC 5370
capLC 3090
CSNK2A1
EC50 [nM]
nLC 692
capLC 776
CSNK2A2
EC50 [nM]
nLC 417
capLC 372
CSNK2B
EC50 [nM]
nLC 513
capLC 537
EIF2AK1
EC50 [nM]
nLC 3311
capLC 3802
EIF2B5
EC50 [nM]
nLC 1549
capLC 21
EIF3J
EC50 [nM]
nLC 525
capLC 550
ERN1
EC50 [nM]
nLC 977
capLC 651
FGFR1
EC50 [nM]
nLC 603
capLC 513
FLT4
EC50 [nM]
nLC 120
capLC 130
GSK3B
EC50 [nM]
nLC 112
capLC 110
IKBKE
EC50 [nM]
nLC 1096
capLC 1122
Relative Response
Concentration [M]

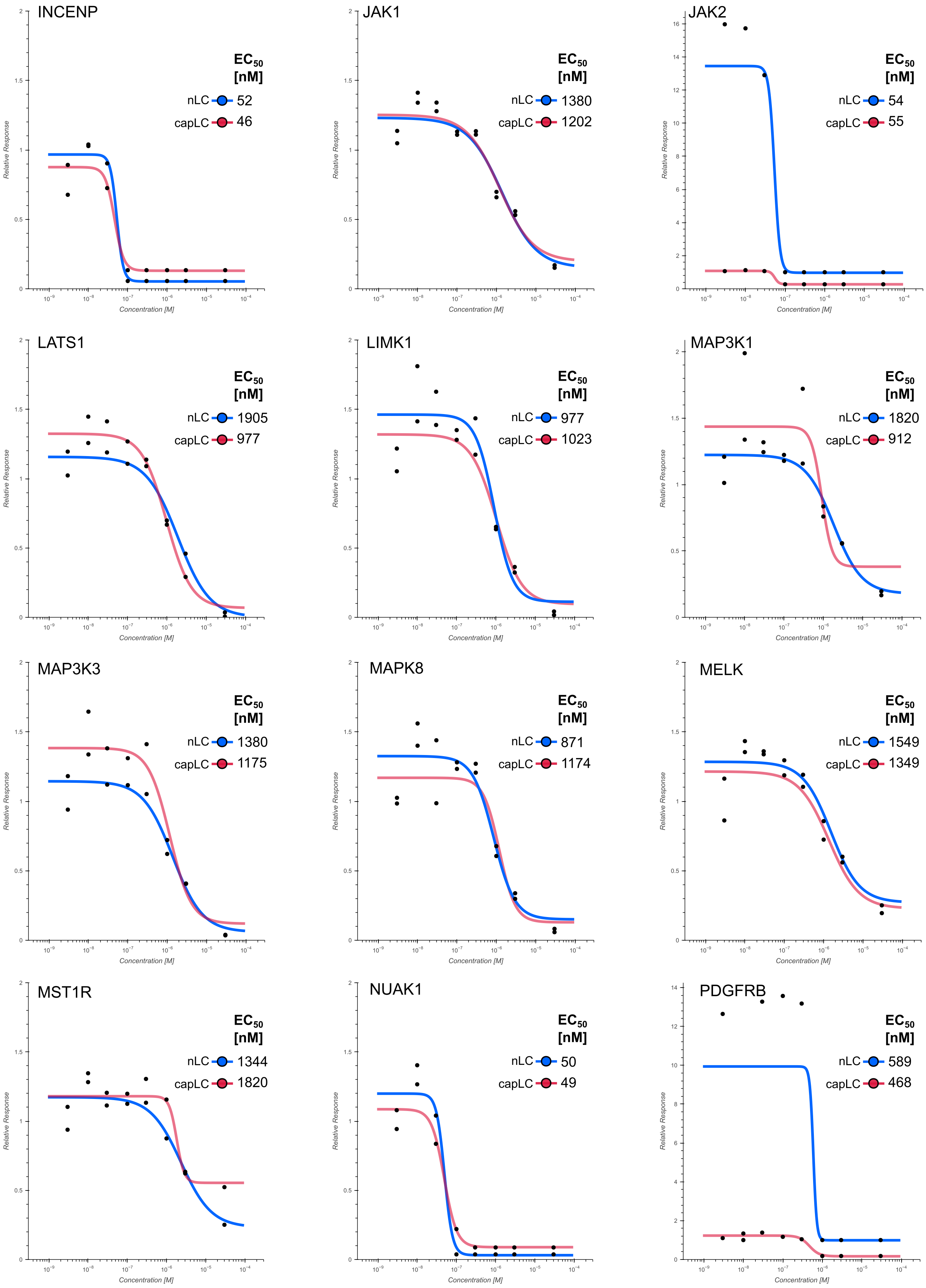
INCENP
EC50 [nM]
nLC 52
capLC 46
JAK1
EC50 [nM]
nLC 1380
capLC 1202
JAK2
EC50 [nM]
nLC 54
capLC 55
LATS1
EC50 [nM]
nLC 1905
capLC 977
LIMK1
EC50 [nM]
nLC 977
capLC 1023
MAP3K1
EC50 [nM]
nLC 1820
capLC 912
MAP3K3
EC50 [nM]
nLC 1380
capLC 1175
MAPK8
EC50 [nM]
nLC 871
capLC 1174
MELK
EC50 [nM]
nLC 1549
capLC 1349
MST1R
EC50 [nM]
nLC 1344
capLC 1820
NUAK1
EC50 [nM]
nLC 50
capLC 49
PDGFRB
EC50 [nM]
nLC 589
capLC 468
Relative Response
Concentration [M]

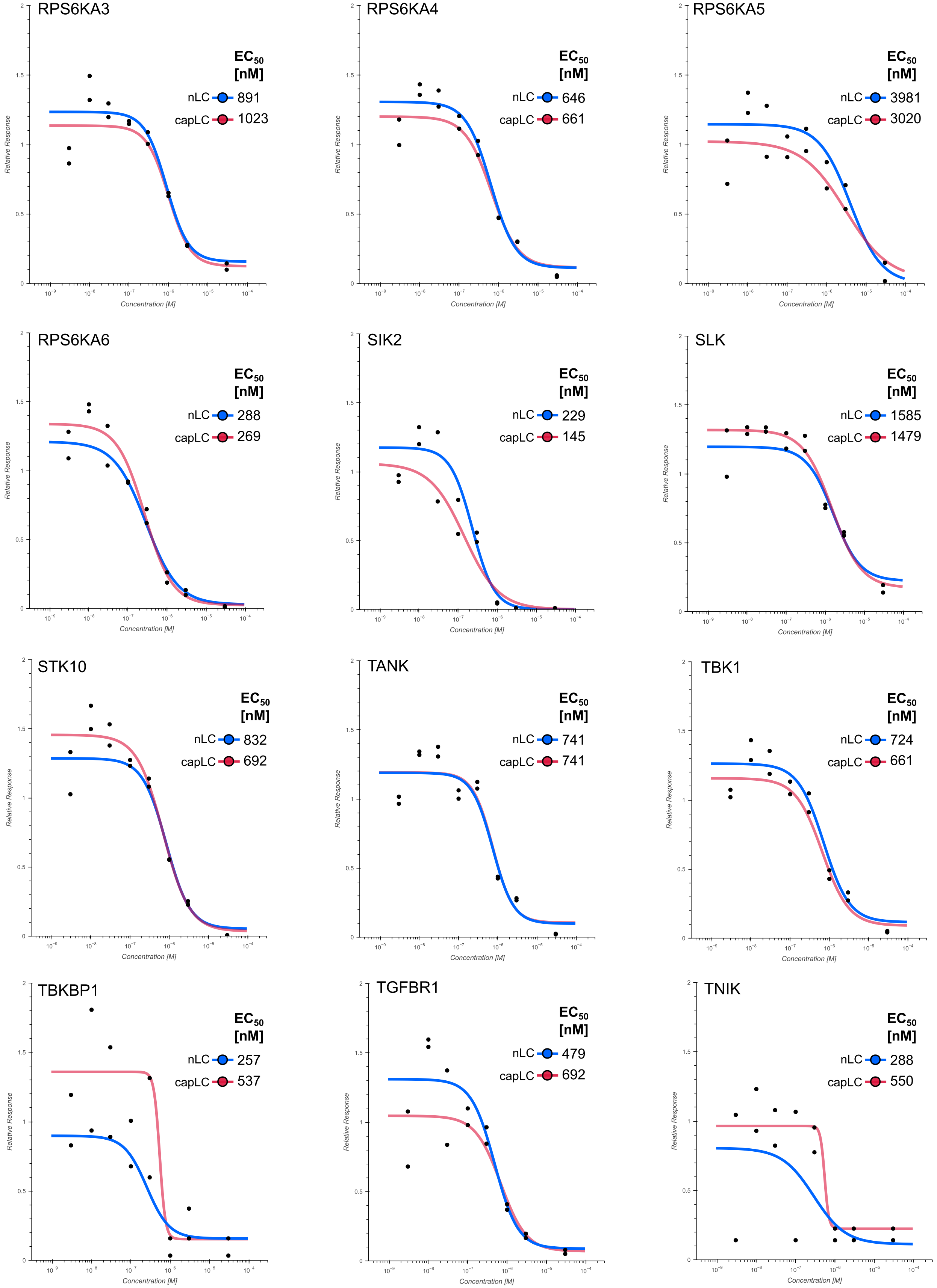

RPS6KA3
EC50 [nM]
nLC 891
capLC 1023
RPS6KA4
EC50 [nM]
nLC 646
capLC 661
RPS6KA5
EC50 [nM]
nLC 3981
capLC 3020
RPS6KA6
EC50 [nM]
nLC 288
capLC 269
SIK2
EC50 [nM]
nLC 229
capLC 145
SLK
EC50 [nM]
nLC 1585
capLC 1479
STK10
EC50 [nM]
nLC 832
capLC 692
TANK
EC50 [nM]
nLC 741
capLC 741
TBK1
EC50 [nM]
nLC 724
capLC 661
TBKBP1
EC50 [nM]
nLC 257
capLC 537
TGFBR1
EC50 [nM]
nLC 479
capLC 692
TNIK
EC50 [nM]
nLC 288
capLC 550
Relative Response
Concentration [M]

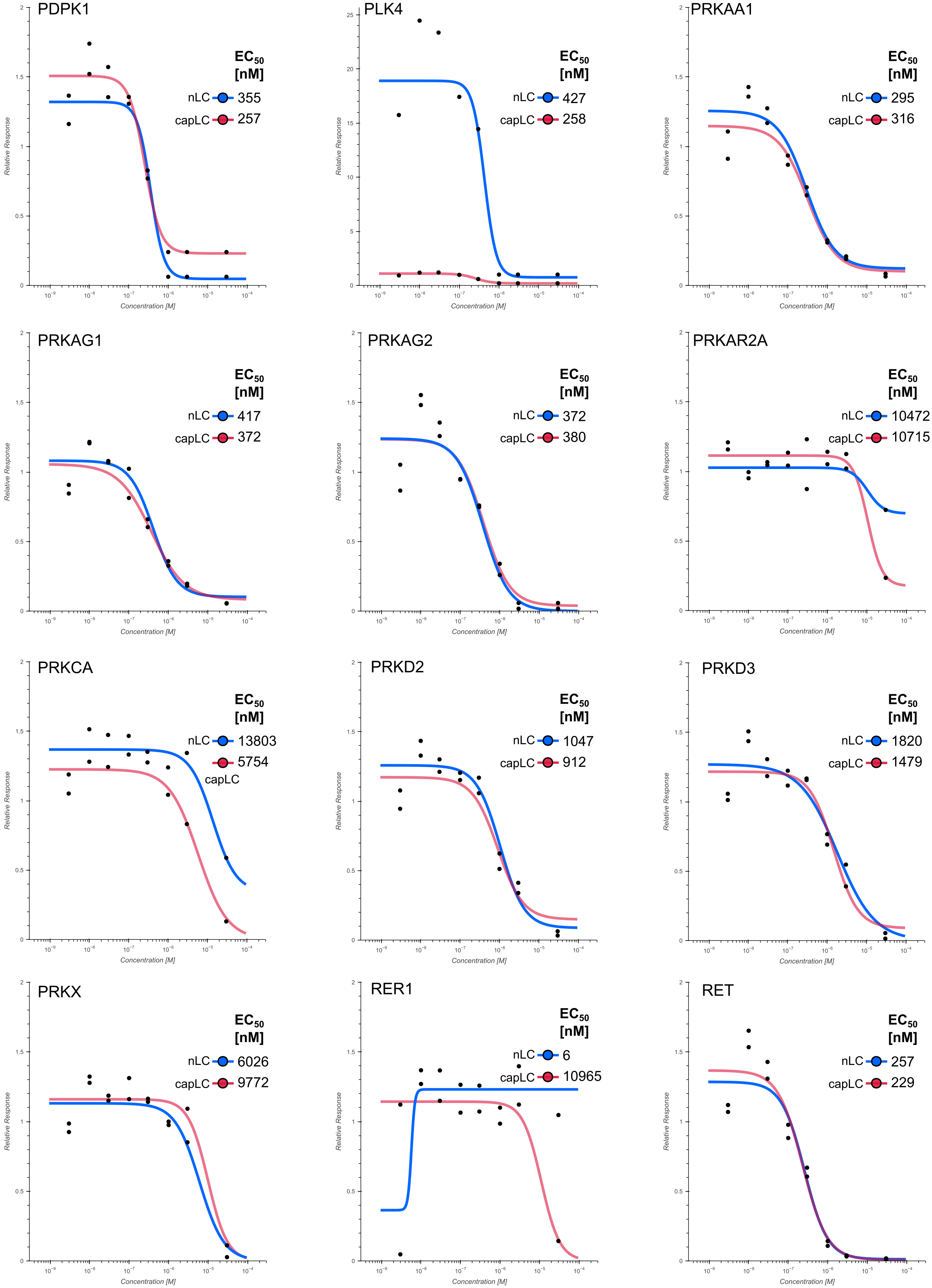
PDPK1
EC50 [nM]
nLC 355
capLC 257
PLK4
EC50 [nM]
nLC 427
capLC 258
PRKAA1
EC50 [nM]
nLC 295
capLC 316
PRKAG1
EC50 [nM]
nLC 417
capLC 372
PRKAG2
EC50 [nM]
nLC 372
capLC 380
PRKAR2A
EC50 [nM]
nLC 10472
capLC 10715
PRKCA
EC50 [nM]
nLC 13803
5754
capLC
PRKD2
EC50 [nM]
nLC 1047
capLC 912
PRKD3
EC50 [nM]
nLC 1820
capLC 1479
PRKX
EC50 [nM]
nLC 6026
capLC 9772
RER1
EC50 [nM]
nLC 6
capLC 10965
RET
EC50 [nM]
nLC 257
capLC 229
Relative Response
Concentration [M]

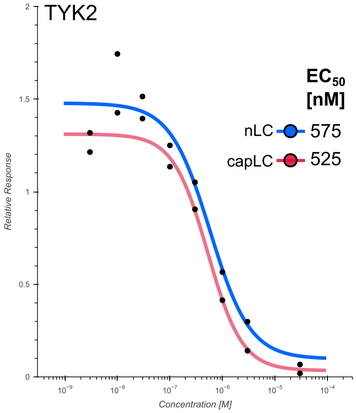
TYK2
EC50 [nM]
nLC 575
capLC 525
Relative Response
Concentration [M]

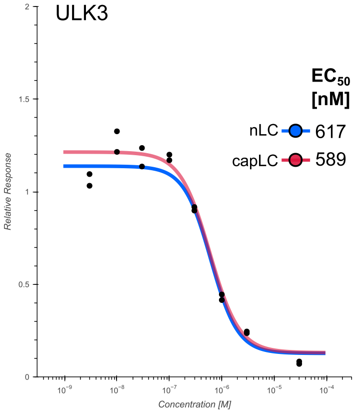
ULK3
EC50 [nM]
nLC 617
capLC 589
Relative Response
Concentration [M]

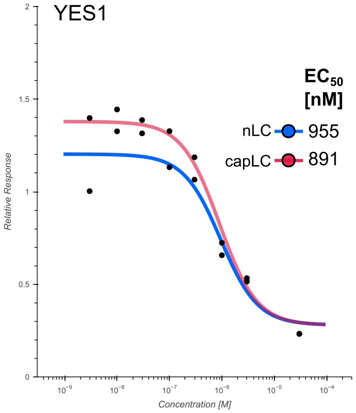
YES1
EC50 [nM]
nLC 955
capLC 891
Relative Response
Concentration [M]

**Supplementary Figure 3.** Dose-response curves and $EC_{50}$ values for all targets of the kinase inhibitor AT-9283 identified by capLC, nLC or both setups.
